# Microfluidic Osteoarthritis-on-a-Chip: Modeling Human Joint Inflammation

**DOI:** 10.64898/2026.02.06.704398

**Authors:** Hosein Mirazi, Scott T. Wood

## Abstract

Osteoarthritis (OA) is a multifactorial joint disease driven by complex interactions among chondrocytes, osteoblasts, fibroblasts, and immune cells across cartilage, bone, and synovial tissues. Conventional monoculture systems are unable to capture this crosstalk, limiting their physiological relevance. Building on our previously established joint-on-a-chip platform, this study evaluated multicellular communication and assessed whether a microfluidic co-culture provides a more realistic representation of joint inflammation compared with monoculture models. Two configurations were established: a healthy, low-inflammation model containing M0 macrophages and an OA-like, high-inflammation model with M1 macrophages. In healthy models, co-culture significantly increased MMP-1 (∼4-fold), MMP-3 (∼15-fold), TIMP-2 (∼5-fold), IL-6 (∼6-fold), and IL-8 (∼5-fold) relative to monoculture, indicating that endogenous signaling initiates basal matrix remodeling and inflammatory pathways. In disease models, M1-driven co-culture elevated MMP-10 (∼300-fold) and MMP-13 (∼60-fold), along with TIMP-2 (∼5-fold), compared with monoculture, reflecting amplified catabolic activation. Direct comparison of disease versus healthy co-culture revealed additional increases in MMP-10 (∼55-fold), MMP-13 (∼95-fold), MCP-1 (∼1.6-fold), MMP-1 (∼1.6-fold), MMP-3 (∼1.8-fold), TIMP-1 (∼1.4-fold), and TIMP-2 (∼1.5-fold), representing a macrophage-mediated shift from homeostasis to OA-like pathology. However, neither IL-1 nor TNFα, each a key inflammatory mediator of OA, differed measurably between healthy and disease models under either monoculture or co-culture conditions. Thus, the microfluidic joint inflammation-on-a-chip model presented here more faithfully recapitulates the pathogenic MMP profile of OA than monoculture systems, but it does not yet fully recapitulate the pathogenic inflammatory environment of OA.

## 2 Introduction

Inflammatory joint diseases, including osteoarthritis (OA), are a leading cause of chronic pain and disability worldwide, collectively affecting over 500 million individuals and imposing a substantial socioeconomic burden upon healthcare systems. [1, 2] OA is characterized by progressive cartilage degradation, subchondral bone remodeling, synovial inflammation, and chronic pain (Figure 1), all of which reduce mobility and quality of life. [3] Despite its increasing prevalence, no disease-modifying OA drugs (DMOADs) have yet been approved for clinical use.[4] Current therapies, such as non-steroidal anti-inflammatory drugs (NSAIDs), corticosteroids, and intra-articular hyaluronic acid injections, offer only short-term symptomatic relief and do not prevent or reverse structural degeneration.[5] This persistent gap highlights the need for physiologically relevant experimental models that can reproduce the complex multicellular interactions along with the inflammatory dynamics of the human joint environment.

**Figure 1.**
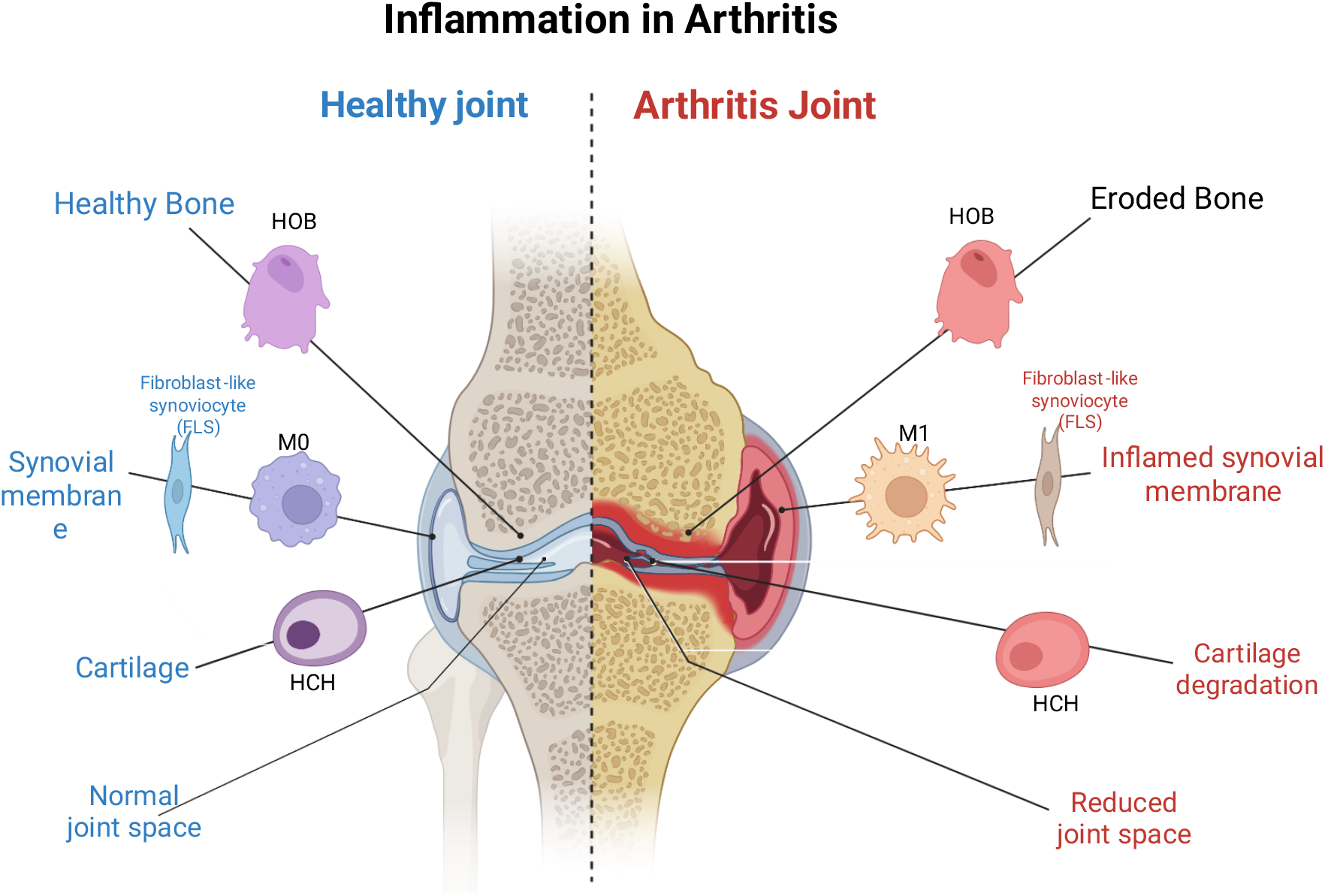
Schematic representation of inflammation in osteoarthritis (OA). Comparison between a healthy joint and an osteoarthritic joint highlighting cell-type-specific roles in joint homeostasis and inflammation. In the healthy joint, homeostatic M0 macrophages coordinate with chondrocytes, osteoblasts, and fibroblast-like synoviocytes to sustain tissue balance and structural integrity. In the OA joint, macrophage polarization toward the pro-inflammatory M1 phenotype drives synovial inflammation, fibroblast activation, cartilage degradation, and bone erosion, leading to narrowing of the joint space and impaired function. Created in BioRender. Mirazi, H. (2025) https://BioRender.com/u04h894

The development of effective OA therapies remains hampered by the limited predictive value of current preclinical models.[6, 7] Traditional *in vitro* monocultures oversimplify the joint microenvironment, failing to capture the paracrine signaling and mechanobiological feedback that drive disease progression.[8] Similarly, animal models, while useful for studying cartilage damage and inflammation, diverge significantly from human joints in cartilage thickness, biomechanical loading, immune regulation, and temporal disease dynamics.[9, 10] These disparities contribute to the frequent translational failure of drug candidates that show early promise but lack efficacy in human trials.[11] In particular, the absence of physiologically integrated *in vitro* models capable of mimicking crosstalk among chondrocytes, osteoblasts, fibroblasts, and immune cells has limited therapeutic discovery and reduced the predictive accuracy of efficacy and safety assessments.[12]

Given these limitations, there is a growing need for human-relevant systems that recapitulate the coordinated biology of the joint. Inflammation within the human joint is not confined to a single tissue; rather, it emerges from complex, coordinated communication among the cartilage, bone, and synovial membrane (Figure 1). These multicellular interactions, mediated by cytokines, growth factors, and extracellular cues, govern whether the joint maintains homeostasis or transitions toward irreversible degeneration (Figure 2D). Conventional monocultures are unable to reproduce these dynamic exchanges, necessitating the creation of integrated human models that can capture joint-level communication and the gradual shift from health to disease.

**Figure 2.**
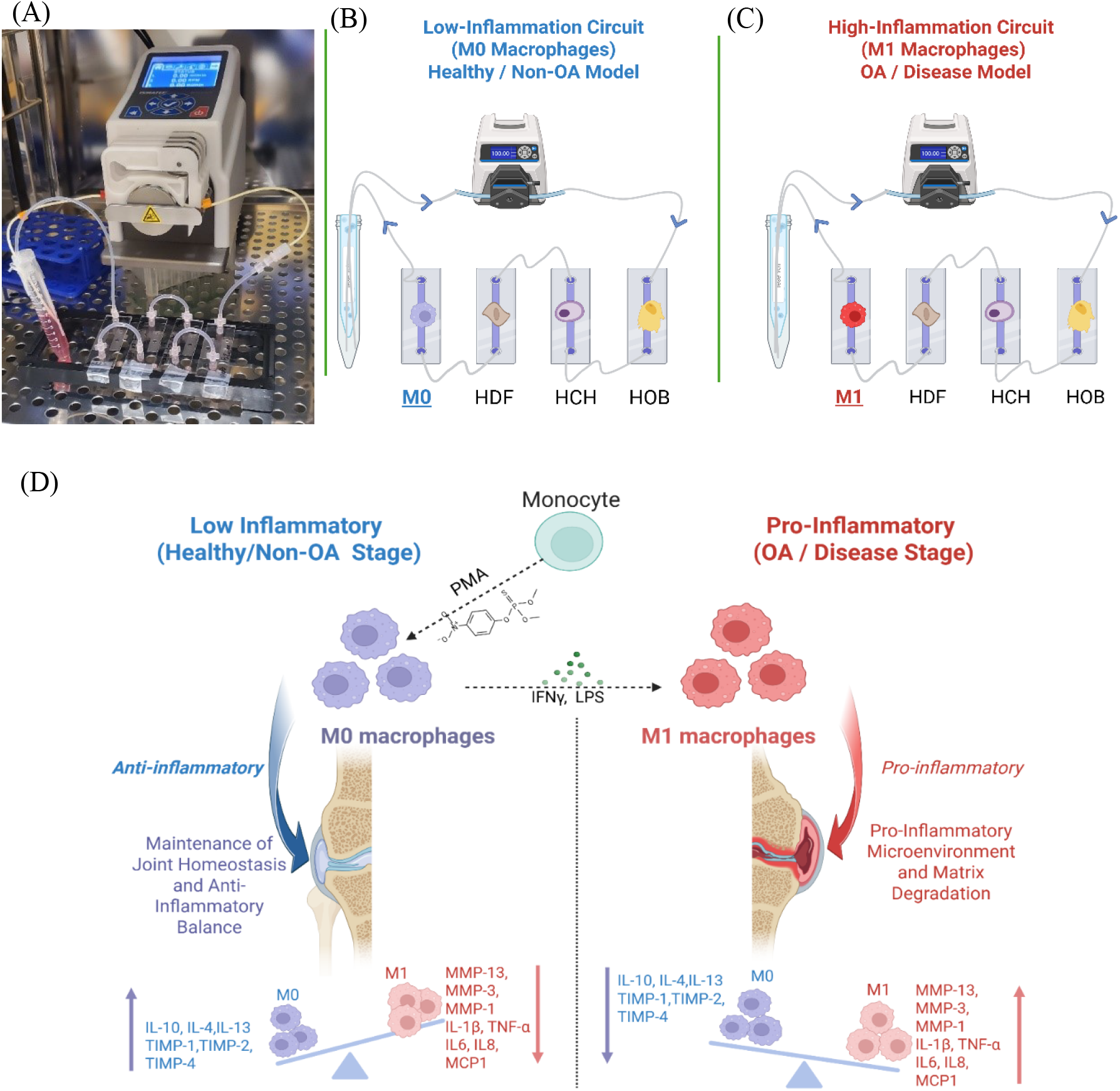
Integrated schematic and experimental representation of the microfluidic joint-on-a-chip platform and macrophage-driven inflammation control. **(A)** Experimental setup showing the connected Ibidi µ-Slide I Luer channels and peristaltic pump used for dynamic recirculation. **(B)** Low-inflammatory (healthy/non-OA) circuit incorporating M0 macrophages, representing physiological joint homeostasis. **(C)** High-inflammatory (OA/disease) circuit incorporating M1 macrophages, modeling a pro-inflammatory microenvironment. **(D)** Schematic of THP-1 macrophage polarization protocol, where monocytes were first treated with phorbol 12-myristate 13-acetate (PMA) to generate M0 macrophages, followed by stimulation with interferon-gamma (IFN-γ) and lipopolysaccharide (LPS) to induce M1 macrophages. The M0 phenotype supports anti-inflammatory balance, while the M1 phenotype promotes joint inflammation and matrix degradation. Created in BioRender. Mirazi, H. (2025)

To address these limitations, we recently presented a microfluidic joint-on-a-chip co-culture system integrating four primary human joint-resident cell types, chondrocytes, osteoblasts, fibroblasts, and macrophages, together within a shared recirculating microenvironment to enable controlled paracrine communication.[13] Two distinct inflammatory states were established in that study and further investigated here: a low-inflammation configuration based on co-culture with quiescent M0 macrophages to model healthy joint conditions, and a high-inflammation configuration based on co-culture with pro-inflammatory M1 macrophages to model generalized inflammatory joint disease, i.e., an ostensibly osteoarthritic (OA)-like disease condition. Unlike conventional exogenous cytokine-stimulated monocultures, this platform leverages macrophage polarization to induce inflammation endogenously, which we expect to generate a more natural, physiologically balanced cytokine profile through immune-stromal crosstalk rather than the more traditional exogenous approaches, which can have unanticipated and unpredictable side effects, especially in co-culture systems.

This study was guided by two central research aims. The first was to quantitatively validate that joint-resident cells engage in functional crosstalk via paracrine signaling within the microfluidic co-culture conditions we previously established. The second was to assess whether the multicellular co-culture system presented here induces more physiologically relevant inflammatory and catabolic responses than conventional monocultures. Together, these objectives provide a framework for evaluating how integrated immune-stromal interactions contribute to the onset and progression of osteoarthritis within a controlled microphysiological environment.

## 3 Materials and Methods

### 3.1 Experimental Design

To evaluate the physiological relevance of the microfluidic joint-on-a-chip platform presented here, two experimental configurations were established: a conventional monoculture and a dynamic microfluidic co-culture, as depicted in Figure 2. The monoculture served as the control for assessing how multicellular communication alters inflammatory and catabolic responses. This parallel design enabled controlled comparison within a fully human *in vitro* context, minimizing variability associated with animal or *ex vivo* models. Within this framework, the present study examined the extent to which the integrated multicellular interactions used here could replicate established inflammatory and remodeling dynamics of OA pathogenesis.

### 3.2 Primary Cell Sources and Culture Conditions

Primary human osteoblasts (HOBs), chondrocytes (HCHs), dermal fibroblasts (HDFs), and THP-1 monocytes were obtained from commercial suppliers (PromoCell and ATCC) and cultured following previously established protocols.[13] Briefly, HOBs, HCHs, and HDFs were maintained in their respective growth media at 37 °C in a humidified atmosphere containing 5 % CO_2_, using passages 2-5 for all adherent cell types. THP-1 monocytes were cultured under standard suspension conditions as described in our prior work.[13]

In the monoculture system, primary human chondrocytes (HCHs) were seeded at 100,000 cells/cm^2^ in 12-well plates (seeding density and surface area comparable to Ibidi µ-Slide channels) and cultured at 37 °C in 5 % CO_2_. Two conditions were established: an untreated control group representing the low-inflammation (i.e., nominally healthy) state and an IL-1β-stimulated group (10 ng/mL; HY-P7028, MedChemExpress, recombinant human, *E. coli*-derived) representing the high-inflammation (disease-like) state. The IL-1β concentration was selected based on prior studies of chondrocyte inflammatory activation.[14, 15] This system served as a reference for evaluating inflammatory responses in the absence of multicellular crosstalk.

In parallel, a microfluidic co-culture model was established to recapitulate joint-level paracrine interactions among HCHs, HOBs, HDFs, and macrophages, as illustrated in Figure 2A-C. For co-culture experiments, each cell type was first expanded independently to ensure stable growth prior to seeding. HCHs were seeded at 80,000 cells/cm^2^, while HOBs and HDFs were each seeded at 40,000 cells/cm^2^ within the microchannels. These densities were selected based on preliminary optimization to achieve approximately 90% confluence for each cell type within the microfluidic channels, ensuring robust attachment and reliable paracrine communication. Macrophages (M0 or M1) were then added to complete the four-cell configuration. A mixed growth medium, also prepared by combining equal volumes of the individual culture media, was used to support metabolic compatibility and paracrine communication among all joint-resident cell types. Also, to preserve cytokine concentrations during the flow phase, the high-inflammation circuit was supplemented with IFN-γ and LPS at the same concentrations used for M1 polarization.

After 3 hours of recirculation, each microfluidic slide was maintained under static conditions for an additional 36 hours to capture post-communication effects. Conditioned media were collected exclusively from the chondrocyte chambers for downstream biomarker analysis. All experiments were performed in biological quadruplicate (n = 4), with identical media compositions and incubation parameters applied to both monoculture and co-culture systems to ensure direct comparability.

### 3.3 Macrophage Differentiation and Polarization

Macrophage phenotypes were generated using a well-established two-step polarization protocol (Figure 2D).[16] Briefly, THP-1 monocytes were seeded at a density of 75,000 cells/cm^2^ in Ibidi µ-Slide I Luer channels and differentiated into M0 macrophages by treatment with 50 nM phorbol 12-myristate 13-acetate (PMA) for 48 hours, followed by a 24-hour rest period in PMA-free medium. These M0 macrophages represented the low-inflammation (healthy) condition. To obtain M1 macrophages for the high-inflammation (OA-like) model, M0 cells were further stimulated for 24 hours with 20 ng/mL interferon-gamma (IFN-γ) and 100 ng/mL lipopolysaccharide (LPS; *E. coli* O111:B4). This sequential activation induced a pro-inflammatory phenotype suitable for integration into the disease-state co-culture model.

### 3.4 Biomarker Quantification via Multiplex ELISA

To evaluate inflammatory signaling and extracellular matrix (ECM) remodeling, conditioned media collected from chondrocyte (HCH) compartments were analyzed using two Quantibody® multiplex sandwich ELISA arrays (RayBiotech, Peachtree Corners, GA, USA). The Human MMP Array Q1 (Cat. #QAH-MMP) quantified MMP-1, -2, -3, -8, -9, -10, -13, and TIMP-1, -2, and -4, while the Human Inflammation Array Q1 (Cat. #QAH-INF) measured IFN-γ, IL-1α, IL-1β, IL-4, IL-6, IL-8 (CXCL8), IL-10, IL-13, MCP-1 (CCL2), and TNF-α. Assays were performed following the manufacturer’s protocol and subsequently submitted to RayBiotech for fluorescence scanning and quantitative analysis. Fluorescence signals were acquired using an Inopsys Innoscan 710AL scanner (Cy3 channel), and data were extracted and normalized using RayBiotech’s analysis software. Relative cytokine and protease levels were compared across experimental groups to assess intercellular signaling and matrix-remodeling activity.

### 3.5 Statistical Analysis

Statistical analyses were performed using GraphPad Prism version 10.3.1 (GraphPad Software, San Diego, CA, USA). Depending on the dataset, either an unpaired two-tailed Welch’s t-test or an ordinary two-way ANOVA followed by Fisher’s least significant difference (LSD) post hoc test with a single pooled variance was applied. Statistical significance was defined as *p* < 0.05. Data are presented as mean ± 95% confidence interval (CI), with each experimental group comprising four biological replicates (*n* = 4).

## 4 Results

As shown in Figure 3, fluorescence-based multiplex ELISA arrays were used to quantify cytokine and matrix remodeling markers in conditioned media collected from chondrocyte compartments. Panels A and B display representative fluorescent scans from the Human Inflammation and Human MMP arrays, respectively, each printed in quadruplicate for precision. Distinct signal profiles were measured in monoculture and co-culture samples, reflecting condition-dependent variations in inflammatory and ECM-remodeling activity. These array outputs formed the basis for subsequent quantitative analyses of individual cytokines, chemokines, metalloproteinases, and their inhibitors.

**Figure 3.**
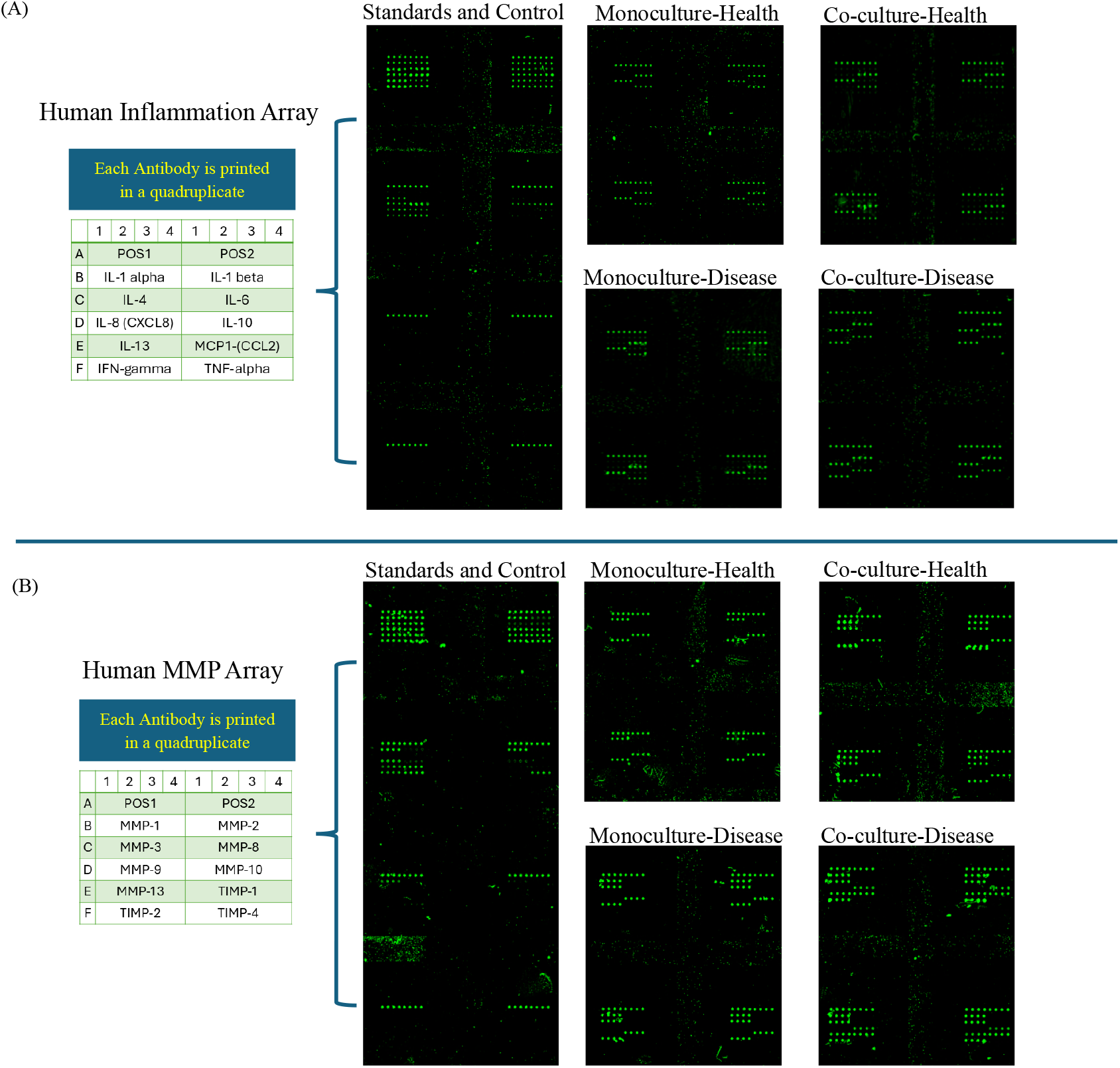
Quantibody® multiplex ELISA arrays used for biomarker quantification across experimental conditions. **(A)** Representative fluorescence scans of the Human Inflammation Array illustrating cytokine expression patterns (IL-1α, IL-1β, IL-4, IL-6, IL-8, IL-10, IL-13, IFN-γ, MCP-1, and TNF-α) across the four experimental conditions: healthy monoculture, disease (IL-1β-stimulated) monoculture, healthy co-culture (M0 macrophages), and disease co-culture (M1 macrophages). **(B)** Representative fluorescence scans of the Human MMP Array showing the relative abundance of matrix remodeling proteins (MMP-1, MMP-2, MMP-3, MMP-8, MMP-9, MMP-10, MMP-13, TIMP-1, TIMP-2, and TIMP-4) under the same four conditions. Each antibody is printed in quadruplicate horizontally to ensure reproducibility and quantitative reliability.

### 4.1 Evaluation of Crosstalk among Joint-Resident Cells

As shown in (Figure 4A), MMP-1 concentrations were markedly elevated in the co-culture compared with the monoculture, increasing from approximately 10 × 10^3^ pg/mL to ≈ 42 × 10^3^ pg/mL (*p* < 0.01), representing an approximate four-fold enhancement. Similarly, under baseline conditions, MMP-3 levels (Figure 4C) were markedly elevated in the co-culture, rising from ≈ 5 × 10^3^ pg/mL to ≈ 80 × 10^3^ pg/mL (*p* < 0.01), corresponding to a roughly 15-fold increase. TIMP-2 (Figure 4I) also increased from ≈ 9.7 × 10^3^ pg/mL in the monoculture to ≈ 48 × 10^3^ pg/mL in the co-culture (*p* < 0.01).

**Figure 4.**
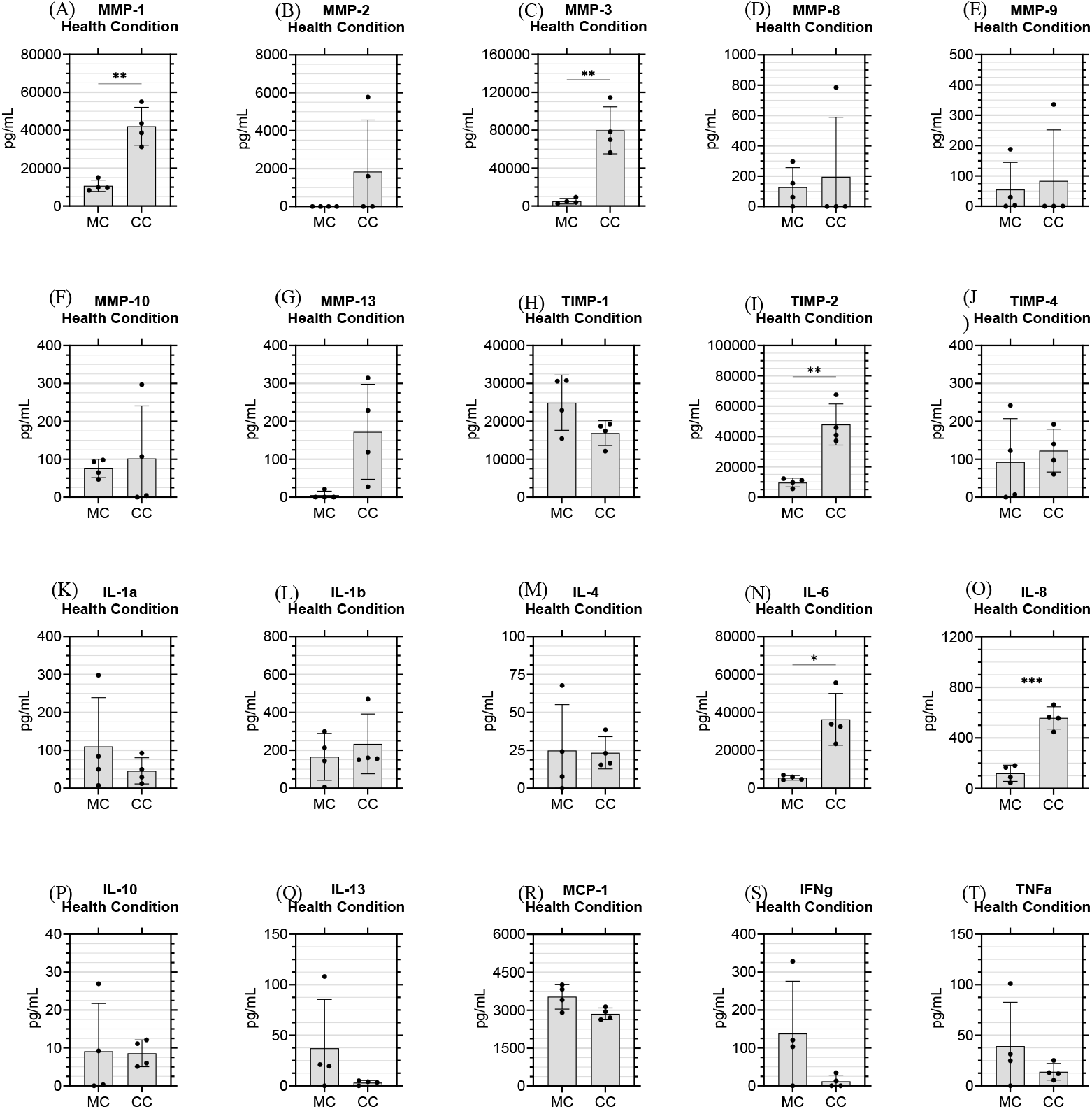
Baseline profiles of matrix-remodeling enzymes and cytokines under healthy (low-inflammation) conditions. Comparison of healthy chondrocyte **monoculture (MC)** and healthy four-cell **co-culture (CC)** containing M0 macrophages. The M0-driven CC exhibited higher secretion of MMP-1, MMP-3, TIMP-2, IL-6, and IL-8, indicating enhanced matrix-remodeling activity and immune-stromal communication even under low-inflammatory conditions. Other markers remained stable between MC and CC, reflecting a balanced, homeostatic joint environment. Data are mean ± 95% CI (n = 4). ^*^p < 0.05, ^**^p < 0.01, ^***^p < 0.001, ^***^p < 0.0001

Among cytokines, IL-6 (Figure 4N) and IL-8 (Figure 4O) had the greatest increases under co-culture conditions. IL-6 levels increased from ≈ 5.5 × 10^3^ pg/mL to ≈ 36 × 10^3^ pg/mL (*p* < 0.05), representing a six-fold increase, while IL-8 secretion rose from ≈ 120 pg/mL to ≈ 560 pg/mL (*p* < 0.001), corresponding to a five-fold enhancement.

Other analytes, including MMP-8 and MMP-9 (Figure 4D, E), remained largely unchanged, whereas MMP-2 and MMP-10 (Figure 4B, F) showed modest increases. MMP-13 (Figure 4G) increased more prominently but with high variability. In contrast, TIMP-1 (Figure 4H) decreased slightly, while TIMP-4 (Figure 4J) exhibited a minor increase. Most cytokines, including IL-1α, IL-1β, TNF-α, IFN-γ, IL-4, IL-10, IL-13, and MCP-1 (Figure 4K-M, P-T), showed minimal variation between conditions, demonstrating a stable low inflammation baseline.

The overall influence of multicellular interactions is illustrated in Figure 5 using a volcano plot to compare co-culture and monoculture for all 20 analytes in a single graph. Here, the x-axis represents the log_2_ fold change (Log_2_FC), where positive values indicate higher expression in the co-culture and negative values denote relative suppression, while the y-axis corresponds to the - log_10_(*p*), reflecting statistical significance. Values above the horizontal dashed line correspond to p < 0.05. In agreement with the marker-specific analyses, IL-8, IL-6, MMP-1, MMP-3, and TIMP-2 clustered in the upper-right quadrant, whereas IL-1α, TNF-α, IFN-γ, IL-13, and MMP-10 appeared in the lower-left quadrant.

**Figure 5.**
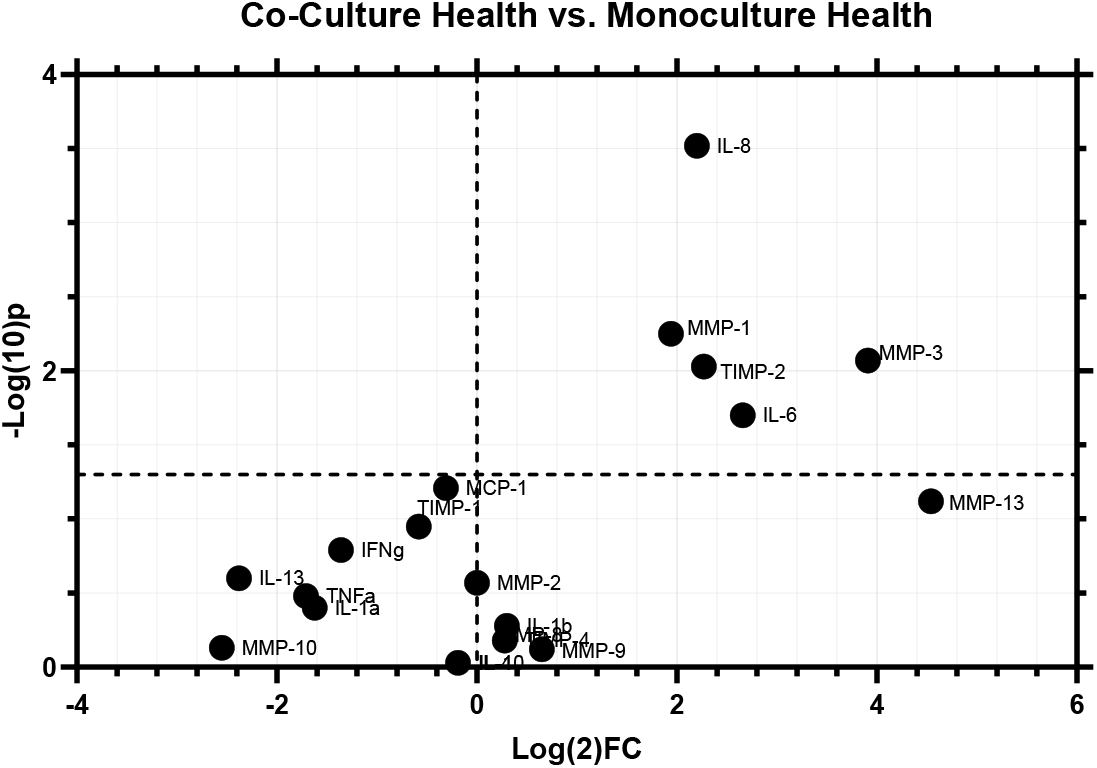
Summary comparison of healthy monoculture and co-culture conditions. The x-axis represents the log_2_ fold change (Log_2_ FC) in analyte expression, where positive values indicate higher levels in the co-culture and negative values indicate higher levels in the monoculture. The y-axis shows the -log_10_(p-value), denoting the statistical significance of each change. Data are presented as mean ± 95 % CI relative to the healthy monoculture.

### 4.2 Inflammatory and Remodeling Responses of Healthy and Disease (OA) Models

As shown in Figure 6A, MMP-1 concentrations increased from approximately 55 × 10^3^ pg/mL in the disease monoculture to ≈ 70 × 10^3^ pg/mL in the disease co-culture (*p* < 0.05). Within the co-culture system, MMP-1 rose from ≈ 42 × 10^3^ pg/mL under healthy (M0) conditions to ≈ 70 × 10^3^ pg/mL in the M1-driven disease state (*p* < 0.001). MMP-3 followed a similar but more pronounced pattern (Figure 6C), increasing from ≈ 81 × 10^3^ pg/mL in the disease monoculture to ≈ 141 × 10^3^ pg/mL in the disease co-culture (*p* < 0.0001), while remaining near 80 × 10^3^ pg/mL under healthy co-culture conditions. MMP-10 (Figure 6F) increased from baseline levels (≈ 75 pg/mL in the healthy monoculture and ≈ 100 pg/mL in the healthy co-culture) to ≈ 5.5 × 10^3^ pg/mL in the M1-driven disease co-culture (*p* < 0.0001). Similarly, MMP-13 (Figure 6G) rose from near-baseline values (≈ 170 pg/mL in the healthy co-culture and ≈ 250 pg/mL in the disease monoculture) to ≈ 17 × 10^3^ pg/mL in the disease co-culture (*p* < 0.0001).

**Figure 6.**
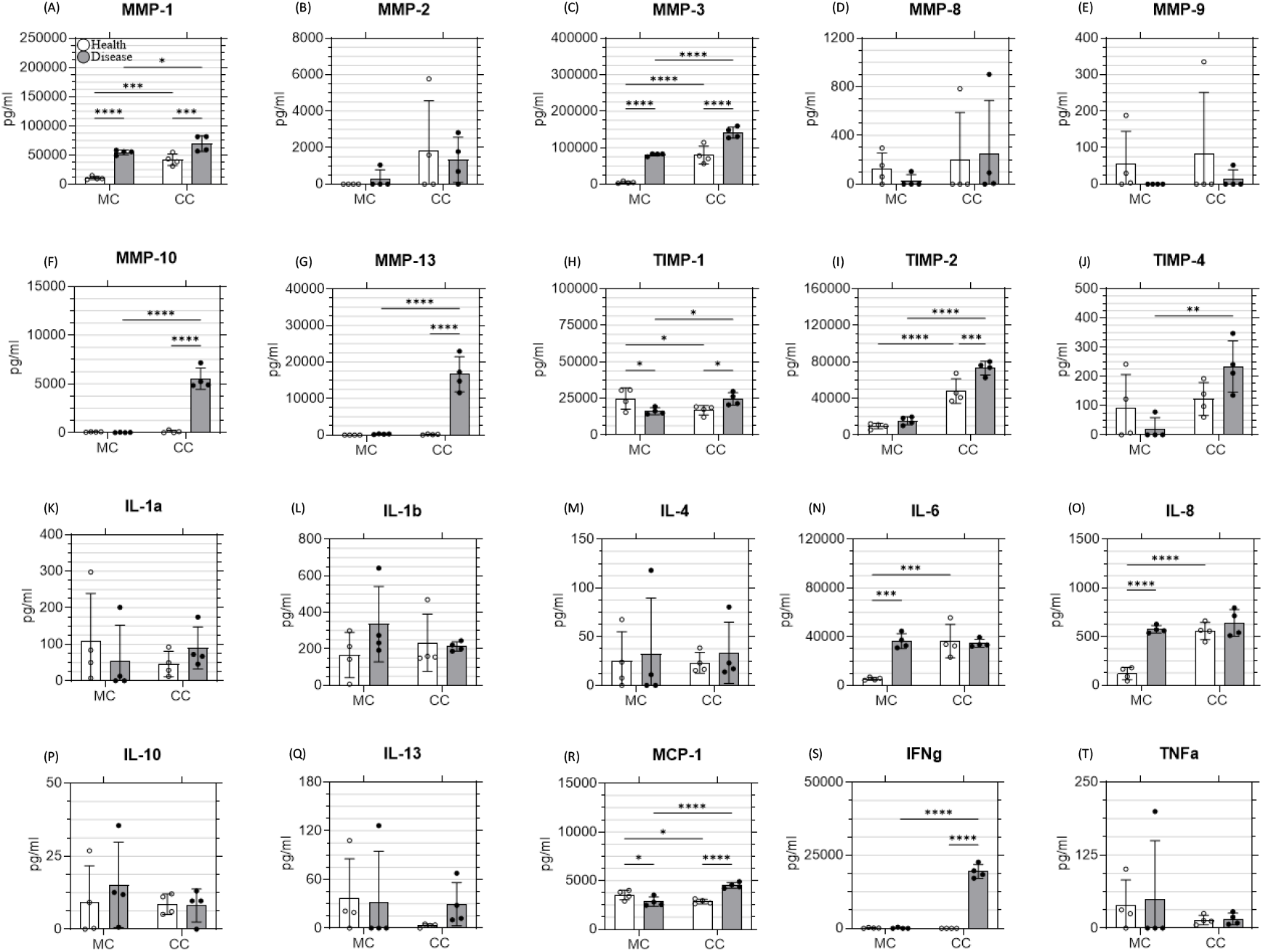
Expression profiles of matrix metalloproteinases (MMPs), tissue inhibitors of metalloproteinases (TIMPs), and cytokines across healthy and OA (disease) conditions in monoculture and co-culture systems. MC denotes chondrocyte **monoculture**, and CC denotes the four-cell **co-culture** (with M0 or M1 macrophages). Panels A-J show MMPs and TIMPs, and panels K-T present ten key inflammatory mediators (IL-1α, IL-1β, IL-4, IL-6, IL-8, IL-10, IL-13, MCP-1, IFN-γ, TNF-α). White bars represent healthy conditions (healthy MC or healthy M0-CC), and gray bars represent OA/disease conditions (IL-1β-stimulated MC or M1-CC). Comparisons were performed within each condition (healthy MC vs healthy CC; disease MC vs disease CC). Data are mean ± 95% CI relative to the corresponding MC. ^*^*p* < 0.05, ^**^*p* < 0.01, ^***^*p* < 0.001, ^****^*p* < 0.0001.

Among the TIMPs, TIMP-1 (Figure 6H) displayed a moderate but significant increase in the disease co-culture (≈ 25 × 10^3^ pg/mL) compared with the disease monoculture (≈ 16 × 10^3^ pg/mL; p < 0.05), while levels in the healthy co-culture remained lower (≈ 17 × 10^3^ pg/mL). TIMP-2 (Figure 6I) showed the strongest elevation, rising from ≈ 15 × 10^3^ pg/mL in the disease monoculture to ≈ 73 × 10^3^ pg/mL in the disease co-culture (*p* < 0.0001), with an intermediate level of ≈ 48 × 10^3^ pg/mL in the healthy co-culture. TIMP-4 (Figure 6J) followed the same pattern, increasing from ≈ 20 pg/mL in the disease monoculture to ≈ 230 pg/mL in the disease co-culture (*p* < 0.01), while remaining at ≈ 122 pg/mL under healthy co-culture conditions.

For cytokine markers, IL-6 (Figure 6N) levels were comparable across all conditions, measuring ≈ 36 × 10^3^ pg/mL in the disease monoculture, ≈ 35 × 10^3^ pg/mL in the disease co-culture, and ≈ 36 × 10^3^ pg/mL in the healthy co-culture. IL-8 (Figure 6O) displayed a similar trend, with concentrations of ≈ 575 pg/mL in the disease monoculture, ≈ 640 pg/mL in the disease co-culture, and ≈ 560 pg/mL in the healthy co-culture. MCP-1 (Figure 6R) levels increased significantly under both inflammatory conditions, rising from ≈ 2.8 × 10^3^ pg/mL in the disease monoculture to ≈ 4.5 × 10^3^ pg/mL in the disease co-culture (*p* < 0.0001). The healthy co-culture maintained lower levels (≈ 2.8 × 10^3^ pg/mL), significantly different from the disease co-culture (*p* < 0.0001). IFN-γ (Figure 6S) showed a striking induction exclusively in the M1 co-culture, reaching ≈ 19.5 × 10^3^ pg/mL, while both the disease monoculture (≈ 100 pg/mL) and the healthy co-culture (≈ 10 pg/mL) remained near baseline (*p* < 0.0001 for both comparisons).

Other analytes, including MMP-2, MMP-8, and MMP-9 (Figure 6B, D, E), showed no significant differences among the four experimental groups. Similarly, cytokines such as IL-1α, IL-1β, IL-4, IL-10, and IL-13 (Figure 6K-M, P, Q, T) remained largely unchanged across all conditions.

To visualize overall biomarker regulation across all experimental groups, volcano plot analyses were performed comparing disease versus healthy states and co-culture versus monoculture systems (Figure 7). The x-axis represents the log_2_ fold change, indicating the magnitude and direction of regulation, while the y-axis denotes the −log_10_(p-value), reflecting statistical significance. In the disease-versus-health comparison (Figure 7A), monocultures (red) displayed modest upregulation of MMP-1, MMP-3, IL-6, and IL-8, consistent with IL-1β stimulation. In contrast, the M1-driven co-culture (blue) exhibited a broader activation profile with higher expression of MMP-10, MMP-13, TIMP-2, IFN-γ, and MCP-1. Similarly, in the co-culture-versus-monoculture comparison (Figure 7B), the healthy (M0) co-culture remained near baseline, whereas the disease (M1) co-culture displayed extensive upregulation, with IFN-γ, MMP-10, and MMP-13 showing the highest fold changes (Log_2_FC ≈ 6-12).

**Figure 7.**
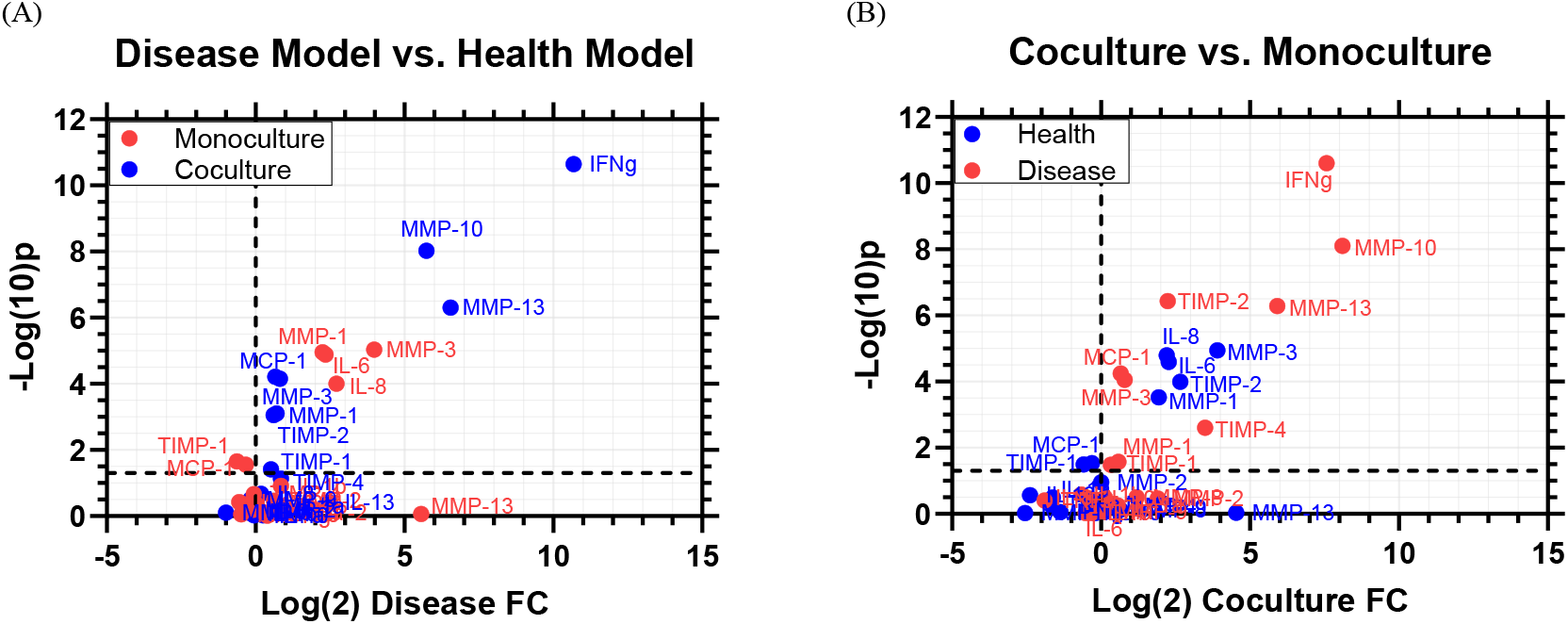
Comparative illustration of global analyte expression across disease, health, and culture configurations. (Left) Log_2_ fold change (x-axis) and -log_10_ p-value (y-axis) comparing disease vs healthy conditions in monoculture (red) and co-culture (blue) systems. (Right) Comparison of co-culture vs monoculture under both health (blue) and disease (red) conditions. Data points represent individual analytes (cytokines, MMPs, and TIMPs). Dashed vertical lines denote no fold-change (Log_2_ FC = 0), and horizontal lines indicate the significance threshold (p = 0.05). These distributions visualize overall inflammatory and matrix-remodeling trends across experimental models.

## 5 Discussion

Despite significant progress in organ-on-a-chip technologies, most osteoarthritis (OA) models remain restricted to dual-cell systems and commonly exclude macrophages cells that are central to synovial inflammation, cytokine signaling, and extracellular matrix (ECM) turnover. Based on the best of our knowledge, no platform has yet fully integrated the four principal human joint-resident cell types chondrocytes, osteoblasts, fibroblasts, and macrophages within a unified, dynamically perfused microenvironment capable of capturing their coordinated interactions.

In this study, we introduced a microfluidic joint-on-a-chip that incorporates all four cell types under shared, recirculating flow to support immune-stromal communication and cartilage-bone coupling. Our work was structured around two central questions: (1) whether joint-resident cells could engage in measurable biochemical crosstalk within a multicellular microfluidic system, and (2) whether this co-culture could reproduce more physiologically representative OA-like inflammatory and catabolic responses compared with conventional monocultures.

To examine how this integrated cellular communication manifests at the molecular level, we next evaluated matrix-remodeling enzymes central to joint homeostasis and degeneration. Matrix metalloproteinases (MMPs) are zinc-dependent endopeptidases that mediate extracellular matrix (ECM) turnover, maintaining normal tissue homeostasis while their dysregulation drives osteoarthritis (OA) and related joint degeneration. Among them, MMP-1, MMP-3, and MMP-13 are principal contributors to cartilage degradation, counterbalanced by tissue inhibitors of metalloproteinases (TIMPs) that preserve ECM integrity and regulate matrix remodeling dynamics. [17, 18]. MMP-1 (interstitial collagenase), responsible for cleaving collagen types I, II, and III, is typically expressed at low levels in healthy cartilage but increases markedly in OA, reflecting its key role in collagen breakdown and joint degeneration. [19, 20]

The strong upregulation of MMP-1 under baseline (non-inflammatory) conditions (Figure 4A) supports the first research question, confirming that functional crosstalk among joint-resident cells is actively maintained within the co-culture. Paracrine communication between chondrocytes, osteoblasts, and fibroblasts further modulated by macrophage-derived mediators appears sufficient to initiate matrix-remodeling activity even without external cytokine stimulation. This coordinated signaling reflects the system’s intrinsic capacity for physiological ECM turnover, positioning MMP-1 as a sensitive indicator of baseline joint function within the microfluidic model.

Beyond MMP-1 driven baseline collagen turnover, the co-culture also modulated broader stromelysin activity through MMP-3. Matrix metalloproteinase-3 (MMP-3, stromelysin-1) degrades multiple extracellular matrix components and can activate other MMPs, amplifying proteolytic remodeling cascades. Although involved in normal tissue repair, it becomes strongly upregulated in osteoarthritic cartilage and synovium, making it a distinguishing biomarker of OA progression. [21-24] Given MMP-3’s broad stromelysin activity and ability to activate other MMPs, its upregulation (Figure 4C) likely reflects macrophage-fibroblast crosstalk that amplifies ECM-regulatory cascades. These results also suggest that the co-culture maintains a dynamically balanced remodeling state even without exogenous inflammatory cues.

TIMP-2 also plays a central role in preserving extracellular matrix (ECM) homeostasis by regulating MMP activity and also contributes to cell differentiation, apoptosis, and cytokine signaling, functioning as an active mediator of tissue remodeling rather than solely an inhibitor. [25-27] The ∼five-fold increase observed in the co-culture relative to monoculture (Figure 4I) likely reflects an adaptive feedback response accompanying MMP upregulation, suggesting that macrophage-driven cues stimulate protective pathways in osteoblasts and fibroblasts. The simultaneous elevation of both MMPs and TIMP-2 might indicate that the co-culture system maintains a controlled remodeling equilibrium, resembling the balanced matrix turnover characteristic of a healthy joint environment.

As a multifunctional cytokine that integrates immune activation, inflammation, and matrix remodeling through gp130 receptor signaling, IL-6 plays a pivotal role in maintaining immune-stromal communication in joint tissues. [28-33]The modest yet reproducible increase observed here (Figure 4N) likely reflects a controlled activation loop inherent to multicellular interactions, rather than overt inflammation. Paracrine signaling between macrophages and fibroblasts may help sustain basal immune vigilance while preserving homeostatic equilibrium. Collectively, these results suggest that IL-6 functions as a sensitive sentinel marker within the four-cell model, maintaining physiological readiness that mirrors the low-grade, regulated cytokine activity characteristic of a healthy joint microenvironment.

In parallel, IL-8 (CXCL8) also exhibited elevated secretion within the co-culture, reinforcing the presence of coordinated paracrine signaling among joint-resident cell types (Figure 4O). As a chemokine that mediates leukocyte recruitment and contributes to cartilage matrix turnover, IL-8 operates through CXCR1 and CXCR2 receptor pathways and is regulated by NF-κB-dependent signaling triggered by cytokines such as TNF-α and IL-1β, as well as mechanical stress. [34-37] The enhancement observed here likely reflects synchronized communication among macrophages, fibroblasts, and chondrocytes that collectively sustain basal immune activity in the microfluidic environment. Rather than indicating pathological inflammation, the elevated IL-8 could represent a controlled state of immune vigilance that preserves homeostatic balance while maintaining responsiveness to inflammatory cues.

Taken together, these results suggest that the four-cell co-culture may support active biochemical crosstalk among joint-resident cells. The coordinated increases in IL-6, IL-8, MMP-1, MMP-3, and TIMP-2—along with the suppression of IL-1α and TNF-α could indicate that the system maintains controlled signaling exchanges rather than remaining quiescent. This pattern might reflect a stable but responsive cytokine-protease network, providing evidence that integrated communication is likely occurring within the co-culture even without external inflammatory stimulation.

Building upon the baseline findings that confirmed functional crosstalk among joint-resident cells, the second research question examined how this multicellular network responds to disease-like (high-inflammation) stimuli and whether it induces more physiologically relevant inflammatory and catabolic responses than conventional monocultures.

The M1-driven co-culture showed selective upregulation (Fig6. A, C, F, G) of MMP-1, MMP-3, MMP-10, and MMP-13, indicating the emergence of coordinated collagenolytic and activation cascades mediated by macrophage-stromal communication. These changes resemble the hierarchical enzyme activation characteristic of osteoarthritic tissue, where stromelysins and collagenases act in concert to remodel the extracellular matrix.

Mechanistically, this enhanced response likely arises from M1 macrophage-derived cytokines, particularly TNF-α and IL-6, which may potentiate fibroblast and chondrocyte activation and sustain a feedback loop consistent with *in vivo* OA pathology. Compared with monocultures exposed to IL-1β alone, the co-culture integrates immune-stromal crosstalk that produces a broader yet regulated inflammatory profile, better reflecting the complex signaling dynamics of the native joint.

Following the observed upregulation of matrix-degrading enzymes, evaluating tissue inhibitors of metalloproteinases (TIMPs) provides critical insight into how the co-culture system maintains homeostatic balance during inflammation-driven remodeling. The substantial rise in TIMP expression under M1-driven conditions (Figure 6H-J) likely reflects engagement of compensatory pathways that preserve ECM integrity, possibly influenced by macrophage-derived cytokines such as IL-10 and TGF-β, which are known to enhance TIMP-2 expression in fibroblasts and chondrocytes.

Regarding the IL and across all three conditions IL-1β-stimulated monoculture, M0 co-culture, and M1 co-culture levels of IL-6 and IL-8 remained comparable (Figure 6N, O) indicating that cytokine release was effectively induced but stabilized within a defined range. The lack of a clear distinction between M0 and M1 macrophage conditions suggests that both phenotypes were capable of sustaining cytokine output at similar magnitudes, possibly reflecting saturation of paracrine feedback mechanisms within the confined microfluidic environment.

Biologically, this plateau could arise from homeostatic regulation, where anti-inflammatory feedback loops limit excessive cytokine accumulation despite ongoing immune activation. Methodologically, it may also reflect signal diffusion constraints or shared downstream receptor sensitivity among joint-resident cells, which collectively moderate cytokine gradients. Rather than diminishing model fidelity, this stability highlights a physiologically relevant low-grade inflammatory equilibrium, representing the steady immune vigilance characteristic of healthy joint tissue. Accordingly, the consistent IL-6 and IL-8 expression across configurations supports the model’s ability to reproduce controlled inflammatory dynamics rather than acute or artifactual activation.

In parallel, MCP-1 (CCL2) further illustrated the macrophage-dependent amplification of inflammatory signaling within the co-culture. The marked elevation observed under M1-driven conditions (Figure 6R) suggests that macrophage polarization enhanced chemotactic activity beyond IL-1β stimulation alone. This trend indicates that macrophage-stromal interactions sustained a feedback loop favoring immune cell recruitment and cytokine propagation, features consistent with *in vivo* osteoarthritic tissue. Biologically, such behavior reinforces the capacity of the microfluidic model to emulate macrophage-mediated recruitment dynamics, positioning MCP-1 as a robust indicator of immune-stromal communication within the OA-like microenvironment.

Also, Interferon-gamma (IFN-γ) functions as a cytokine that bridges immune activation and tissue catabolism within the joint microenvironment. It amplifies inflammation by inducing TNF-α, IL-6, and MMP-13 through STAT1- and PKR-dependent pathways, thereby coordinating immune activation with extracellular-matrix degradation. Through these mechanisms, IFN-γ may act as a key amplifier of osteoarthritic progression, sustaining macrophage-stromal activation and promoting degenerative remodeling of joint tissues. [38]

Similarly, IFN-γ showed a distinct induction confined to the M1 co-culture (Figure 6S), underscoring that macrophage polarization rather than IL-1β stimulation was the dominant driver of immune activation. This response highlights the establishment of a macrophage-dominated inflammatory axis capable of reinforcing cytokine and matrix-degrading cascades within the joint microenvironment. While part of this elevation may stem from residual or feedback-amplified IFN-γ introduced during polarization, the overall trend reflects genuine immune-stromal crosstalk characteristic of chronic osteoarthritic inflammation.

The volcano plots (Figure 7) further reinforce these findings by showing broader activation in the M1-driven co-culture than in the IL-1β-stimulated monoculture. The co-culture displayed coordinated increases in catabolic enzymes (MMP-10, MMP-13) and regulatory mediators (TIMP-2, MCP-1, IFN-γ), indicating macrophage-dependent immune-stromal signaling. In contrast, the monoculture response was narrower and predominantly cytokine driven.

Collectively, these integrated patterns support the second research question by indicating that the four-cell co-culture may elicit a more balanced yet multifaceted OA-like response than monocultures. Rather than producing a narrow cytokine-driven activation profile, the co-culture appears to sustain an inflammatory state that bridges homeostatic and disease-like conditions, reflecting a broader and more physiologically relevant joint response.

## 6 Conclusions

This study presents a four-cell human joint-on-a-chip model that reproduces key features of osteoarthritis (OA), including macrophage-driven inflammation and extracellular matrix (ECM) remodeling. By integrating chondrocytes, osteoblasts, fibroblasts, and macrophages within a shared microenvironment, the platform captures essential immune-stromal crosstalk and more closely reflects human joint physiology than conventional monocultures.

A central limitation of the current system is the use of IFN-γ and LPS for macrophage polarization, which more closely model bacterial activation than sterile OA-associated inflammation. Future iterations could incorporate disease-relevant cues such as IL-1β, TNF-α, or OA synovial fluid to generate more physiologically representative macrophage phenotypes. Additional expansion to include fibroblast-like synoviocytes (FLS), T cells, or other immune populations may further enhance biological complexity and predictive value.

Despite these limitations, the platform demonstrates strong potential for preclinical evaluation of disease-modifying OA drugs (DMOADs) and targeted anti-inflammatory compounds. Its ability to capture both degradative and regulatory signaling under controlled, human-relevant conditions positions this model as a promising translational tool for studying joint inflammation and therapeutic modulation.

Importantly, although the disease co-culture exhibited marked upregulation of several inflammatory markers, the unstimulated healthy co-culture may still serve as the more reliable predictive model. Despite the strong inflammatory activation seen in the disease co-culture, the healthy co-culture may remain the more reliable predictor of therapeutic response, as its behavior reflects the natural signaling capacity of M0 macrophages without artificial polarization cues. This likely reflects the inherent ability of M0 macrophages to sustain endogenous, low-level signaling that more closely mirrors early-stage OA physiology than the exaggerated IFN-γ/LPS-induced inflammatory response.

## Acknowledgments

This material is based upon work supported by the National Science Foundation under Grant No. 2517512 (2234590).

## Ethics Statement

Primary human osteoblasts, chondrocytes, fibroblasts, and macrophages used in this study were obtained from commercial sources (PromoCell and ATCC). All cell lines were anonymized and supplied with verified informed consent and ethical clearance from the vendors; therefore, no additional institutional ethical approval was required for this work.

## Use of Generative Artificial Intelligence

A generative artificial intelligence (AI) tool was used in a limited capacity to enhance language quality and readability during the preparation of this manuscript. All scientific content, data interpretation, experimental design, and conclusions were independently conceived, developed, and rigorously verified by the authors. The authors retain full responsibility for the accuracy, integrity, and originality of the work.

## Author Contributions

HM: Conceptualization, Data Curation, Formal Analysis, Investigation, Methodology, Software, Validation, Visualization, Writing – original draft, Writing – review and editing. SW: Conceptualization, Data Curation, Formal Analysis, Funding Acquisition, Investigation, Methodology, Project Administration, Resources, Software, Supervision, Validation, Visualization, Writing – original draft, Writing – review and editing.

